# Individual alpha peak frequency predicts 10 Hz flicker effects on selective attention

**DOI:** 10.1101/185132

**Authors:** Rasa Gulbinaite, Tara van Viegen, Martijn Wieling, Michael X Cohen, Rufin VanRullen

## Abstract

Rhythmic visual stimulation (“flicker”) is primarily used to “tag” processing of low-level visual and high-level cognitive phenomena. However, preliminary evidence suggests that flicker may also entrain endogenous brain oscillations, thereby modulating cognitive processes supported by those brain rhythms. Here we tested the interaction between 10 Hz flicker and endogenous alpha-band (~10 Hz) oscillations during a selective visuospatial attention task. We recorded EEG from human participants (both genders) while they performed a modified Eriksen flanker task in which distractors and targets flickered within (10 Hz) or outside (7.5 or 15 Hz) the alpha band. By using a combination of EEG source separation, time-frequency, and single-trial linear mixed effects modeling, we demonstrate that 10 Hz flicker interfered with stimulus processing more on incongruent than congruent trials (high vs. low selective attention demands). Crucially, the effect of 10 Hz flicker on task performance was predicted by the distance between 10 Hz and individual alpha peak frequency (estimated during the task). Finally, the flicker effect on task performance was more strongly predicted by EEG flicker responses during stimulus processing than during preparation for the upcoming stimulus, suggesting that 10 Hz flicker interfered more with reactive than proactive selective attention. These findings are consistent with our hypothesis that visual flicker entrained endogenous alpha-band networks, which in turn impaired task performance. Our findings also provide novel evidence for frequency-dependent exogenous modulation of cognition that is determined by the correspondence between the exogenous flicker frequency and the endogenous brain rhythms.

**Significance:** Here we provide novel evidence that the interaction between exogenous rhythmic visual stimulation and endogenous brain rhythms can have frequency-specific behavioral effects. We show that alpha-band (10 Hz) flicker impairs stimulus processing in a selective attention task when the stimulus flicker rate matches individual alpha peak frequency. The effect of sensory flicker on task performance was stronger when selective attention demands were high, and was stronger during stimulus processing and response selection compared to the pre-stimulus anticipatory period. These findings provide novel evidence that frequency-specific sensory flicker affects online attentional processing, and also demonstrate that the correspondence between exogenous and endogenous rhythms is an overlooked prerequisite when testing for frequency-specific cognitive effects of flicker.

## INTRODUCTION

A popular technique to study selective attention (spatial, feature-based, object-based) relies on brain responses to rhythmic sensory stimulation (visual flicker, amplitude-modulated sound, or tactile vibrations). In M/EEG recordings, responses to rhythmic stimuli are periodic, with differentiable spectral signatures at frequencies identical or harmonically related to the stimulus (“steady-state visual evoked potentials”; SSVEPs). The frequency of SSVEPs is determined by the stimulus and is stable over time (Regan, 1966; Herrmann, 2001; Keitel et al., 2017), whereas the amplitude is time-varying and depends on cognitive variables, including attention (Morgan et al., 1996). SSVEP amplitude is higher for attended vs. ignored stimuli, making the SSVEP an important tool for measuring selective attention continuously over space and time (Regan and Heron, 1969; coined "frequency tagging" by Tononi et al., 1998).

There is an implicit assumption underlying the frequency tagging approach: The stimulus rhythm does not interact with ongoing endogenous brain rhythms, and therefore the choice of tagging frequency, according to some, should not have behavioral consequences (Keitel et al., 2014). In other words, frequency tagging allows measuring cognitive processes without influencing those processes. However, this assumption is difficult to reconcile with empirical evidence of stronger SSVEPs when periodic light flashes are “in-sync” with individual alpha peak frequency (IAF; Adrian and Matthews, 1934; Notbohm et al., 2016). The tagging frequency neutrality assumption is also inconsistent with differences in SSVEP amplitude across brain areas and stimulation frequencies. For example, visual areas show strong SSVEPs to alpha and gamma flicker (Regan, 1989), frontal areas respond strongly to theta (4-8 Hz; Srinivasan et al., 2007), and SSVEP to face stimuli are most pronounced at rates ~6 Hz (Alonso-Prieto et al., 2013).

These observations suggest an alternative hypothesis: Exogenous rhythmic stimulation, at least at certain frequencies, can entrain endogenous brain rhythms, and *modulate* cognitive processes supported by those brain rhythms (Mathewson et al., 2012; de Graaf et al., 2013; Spaak et al., 2014). A critical test for the idea of frequency-specific behavioral effects of flicker, however, requires demonstrating that stimulation rates close to a “natural” frequency of the network engaged in the task have the strongest effect on behavior.

Previous studies assessed the behavioral effect of alpha-band rhythmic visual stimuli, and provided exciting but limited evidence due to the following reasons. First, the relationship between flicker and IAF, and variability of IAF across brain regions (Haegens et al., 2014; Gulbinaite et al., 2017) were not taken into account. Second, most studies used one or two flicker frequencies (Mathewson et al., 2012; Spaak et al., 2014; Kizuk and Mathewson, 2017) (but see de Graaf et al., 2013), preventing conclusions about frequency-band specific behavioral effects. Third, because the effects of alpha-band flicker were assessed *after* stimulation train offset (Mathewson et al., 2012; de Graaf et al., 2013; Spaak et al., 2014), the effects could have been due to temporal expectations induced by rhythmic stimulation, as opposed to entrainment of endogenous oscillations (Breska and Deouell, 2014).

Here we tested the effect of alpha flicker during stimulus processing in a visuospatial attention task. Modulation of alpha-band oscillations is observed both in preparation to the upcoming stimulus (Frey et al., 2015), and during stimulus processing (van Diepen et al., 2016). Thus, we hypothesized that alpha flicker could interfere with functioning of networks operating in alpha and, as a consequence, interfere with selective attention. We used the Eriksen flanker task (Gulbinaite et al., 2014), with target and flankers flickering within or outside the alpha band. Based on current theories of the inhibitory role of alpha-band oscillations (Jensen and Mazaheri, 2010; Klimesch, 2012), and retinotopic mapping of SSVEPs (Di Russo et al., 2007; Cottereau et al., 2011), we reasoned that stimulus processing in parts of the visual field flickering in alpha will be impaired. We found, using mixed-effects regression modeling of single-trial time-frequency and SSVEP responses, that inhibitory effects of alpha flicker on task performance were most pronounced when alpha flicker frequency matched the IAF peak.

## METHODS

### Participants

Thirty-one participants were recruited using the online participant recruitment system of the Psychology Department at the University of Amsterdam, and took part in the experiment in exchange for a course credit or monetary compensation (€15). Participants with a first-degree family member with epilepsy or migraine were excluded from this study. Participants had normal or corrected-to-normal vision, no reported history of psychiatric disorders, and were self-reported right-handed. The experiment was approved by the local ethics committee of the University of Amsterdam and informed consent was obtained from all participants. Data from four participants were excluded from the analyses: One due to technical issues during EEG recording, and three due to poor behavioral performance (accuracy < 80%). Thus, the final sample was twenty-seven participants (15 female, mean age 22.4).

### Stimuli and procedure

A modified version of the Eriksen flanker task with a four-to-two mapping of stimuli to response buttons was used (Wendt et al., 2007; Gulbinaite et al., 2014). The experimental paradigm was similar to that described in the study by Gulbinaite et al. (2014), in which participants were instructed to respond to the central target letter and ignore the surrounding distracting flanker letters. Instead of the previously employed linear configuration of target and flankers, here flanker stimuli were equidistantly positioned relative to the centrally presented target stimulus (Fig. 1). This was done to maximize the SSVEP signal related to the processing of the flankers (Vanegas et al., 2013). For the very same reason, each letter stimulus size was increased to ~3.35° of visual angle, separated by ~0.65° visual angle. The total estimated cortical representation of the stimuli in V1 was 77.09 and 97.72 (arbitrary units) for targets and flankers respectively (using cortical magnification factors provided by Schira et al., 2007). SSVEPs were elicited by modulating the luminance of stimuli by a 7.5 Hz, 10 Hz, or 15 Hz sine wave (Fig. 1B). Sine-wave stimulus luminance modulation allows greater frequency precision (energy in the stimulus at the harmonic frequencies is negligible as compared to the fundamental stimulation frequency), and also ensures equal total luminance for different frequencies within a given time window of flicker. The stimuli were presented in Sloan font (Pelli et al., 1988), letters of which are equally discriminable and for which height equals width. Stimuli were displayed on a 23-in. LCD monitor with a resolution of 1920 × 1080 pixels and a refresh rate of 120 Hz. The viewing distance was unconstrained and kept at approximately 90 cm.

**Figure 1.**
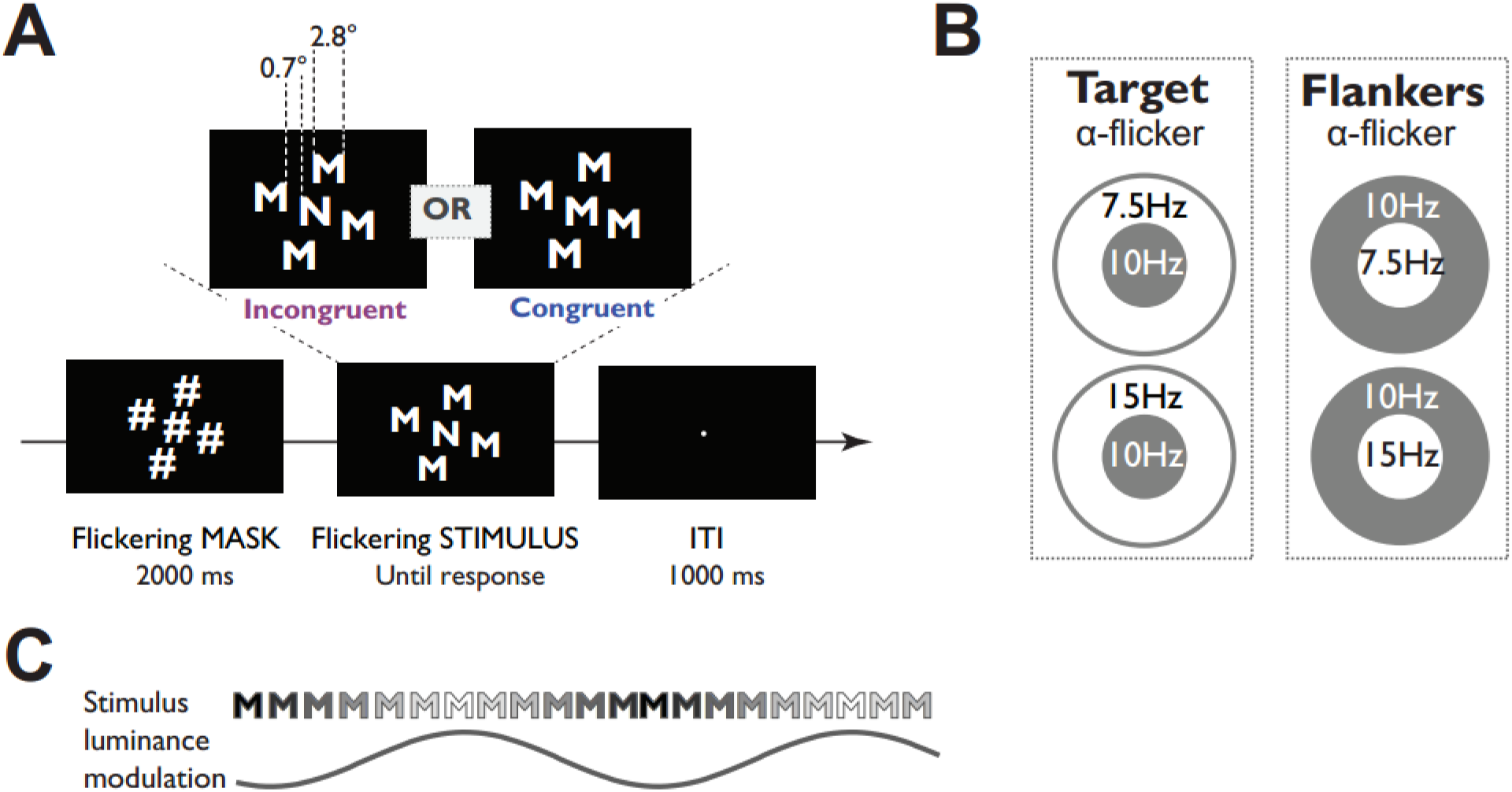
Stimuli and task. (A) Trials began with the presentation of a mask consisting of hash marks for 2000 ms followed by imperative stimulus, presentation of which lasted until a button press or until the deadline of 1200 ms was exceeded, and was followed by an inter-trial interval (ITI) of 1000 ms. (B) Each stimulus consisted of a target letter flickering within (10 Hz) or outside (7.5 or 15 Hz) the alpha-band range, while flanker letters flickered outside (7.5 or 15 Hz) or within alpha-band range respectively. (C) Frequency tagging was implemented by sine-wave modulation of stimulus luminance.

Stimuli consisted of a target letter and four identical flanker letters presented in white against a black background. Participants used a computer mouse to respond. Four letters (E, F, M, N) were used as stimuli and were mapped onto two response keys: When the central target letter was M or E participants pressed the left button with their left thumb, and when the central target was N or F participants pressed the right button with their right thumb. Only response congruent (e.g., M M M M M) and response incongruent (e.g., M M N M M) stimuli were used in the experiment. The order of different types of trials was pseudo-randomized, with the constraint that exact stimulus-response repetitions were not presented; this prevents repetition priming effects (Mayr et al., 2003). The overall probability of congruent and incongruent trials, as well as the proportion of left-and right-hand responses, was kept equal. Participants completed a practice session (40 trials over 4 blocks), which was followed by the experiment session (12 blocks, with 56 trials per block). In the practice session, feedback on performance was given after each trial; in the experiment session, average performance feedback was given after each block.

Each trial started with a 2 s presentation of a pre-stimulus mask comprising hash marks. The hash marks in the four distractor positions flickered at a common frequency, while the hash mark in the central target position flickered at a different frequency. All trials included 10 Hz flicker, thus creating 4 tagging conditions in total (Fig. 1C). The purpose of the 10 Hz flicker was to test our key hypothesis about entrainment effects of alpha flicker on endogenous alpha-band neural oscillations. 7.5 Hz and 15 Hz tagging frequencies were selected to be outside the typical alpha-band range while still being perceptually similar to the 10 Hz flicker.

Despite growing evidence for the entrainment of endogenous oscillations by rhythmic stimuli (Spaak et al., 2014; Notbohm et al., 2016), previously reported behavioral effects of 10 Hz rhythmic stimulation on perception and attention can alternatively be explained by temporal expectations because the imperative stimulus was presented after rhythmic stimulation offset (Mathewson et al., 2012; de Graaf et al., 2013; Spaak et al., 2014). The issue of entrainment vs. temporal expectation is a subtle but important distinction, particularly considering the involvement of alpha-band oscillations in temporal expectation (Rohenkohl and Nobre, 2011). Although arguments have been made against this potential alternative (Spaak et al., 2014), our design avoids the issue by having the imperative stimulus flickering all the time.

Tagging frequencies were changed on each block, and blocks were randomly presented to participants. After 2 s the mask was replaced by the imperative stimulus (e.g. N N M N N) flickering at the same frequencies as the hash marks they replaced. Stimulus presentation lasted until a response was made, or until the deadline of 1200 ms after the stimulus onset was exceeded. Participants were instructed to respond as quickly and as accurately as possible. In order to prevent eye movements, participants were instructed to focus on a small green dot presented continuously at the center of the screen.

### Data acquisition and preprocessing

EEG data were acquired at 1024 Hz using a 64-channel BioSemi system (http://www.biosemi.com). Two additional electrodes were placed on the outer eye canthi to record horizontal eye movements (re-referenced offline to a single bipolar channel). Data analyses were performed using EEGLAB and custom written Matlab scripts. Offline the data were high-pass filtered at 0.5 Hz, and re-referenced to the average of all scalp electrodes. Thereafter, the data were epoched (-1.5 to 3.5 s relative to the mask onset, which is sufficiently long to prevent potential edge artifacts from contaminating the analysis windows), and baseline-corrected with respect to the time window of −200-0 ms (where 0 corresponds to mask onset). Bad electrodes, trials containing muscle artifacts (during mask and imperative stimulus presentation period), as well as trials containing horizontal eye movements away from fixation, and trials containing eye blinks during imperative stimulus presentation were manually rejected. Trials containing single-blinks (but not many successive blinks) during mask period – on average, 4.12% (SD = 5.29%) – were not removed, as the spatiotemporal filtering method used to extract SSVEP responses suppresses blink-related artifacts. Next, we performed independent component analysis (ICA) using the JADE algorithm (Delorme and Makeig, 2004). Independent components (ICs) easily detectable as being driven by eye movements, EMG, or noise were identified (following criteria provided by Chaumon et al., 2015). These artifactual ICs (average of 1.85 ICs per participant) were removed only for analyses involving identification of IAF and electrode-level theta time-frequency power. Analyses involving spatiotemporal source-separation (described in the following section) did not require removing artifactual independent components and were done on the full-rank data.

Trials with no or incorrect responses, as well as trials with reaction times faster than 150 ms or 3 standard deviations above the mean in each condition were excluded from the analysis. The average number of trials per condition included in the statistical analysis for both EEG and behavioral data was 278 (SD = 22.15) for congruent trials, and 269 (SD = 21) for incongruent trials.

### SSVEP analyses: spatiotemporal filtering

Frequency- and stimulus-specific (target vs. flankers) SSVEP responses were obtained using a spatiotemporal source separation method called rhythmic entrainment source separation (RESS), which combines both temporal (flicker frequency) and spatial (topographical distribution) characteristics of SSVEPs (Cohen and Gulbinaite, 2017). The RESS method determines an optimal spatial filter (electrode weights) that maximally separates frequency-specific SSVEPs (the signal, S) from non-SSVEP brain activity (the reference, R). We used brain activity at neighboring frequencies as reference data. Thus, instead of analyzing SSVEPs from channels with maximum power at the tagging frequency, we analyzed a linearly weighted combination of all electrodes. In addition to increasing the SSVEP signal-to-noise ratio, RESS also helps separate the SSVEP-related activity from temporally cooccurring non-SSVEP-related activity such as blinks, stimulus evoked responses, and activity at other frequencies (Cohen and Gulbinaite, 2017).

For each participant, six spatial filters were constructed (separately for each tagging frequency, and stimulus type) because: (1) SSVEP topographies differ for centrally presented target and peripheral flankers (Fig. 2); (2) different frequency SSVEPs have different sources and therefore different scalp projections (Heinrichs-Graham and Wilson, 2012; Lithari et al., 2016); (3) SSVEP topographies may differ across participants due to anatomical differences. Each spatial filter was designed as follows. First, condition-specific single-trial data were concatenated and temporally filtered using three different narrow-band Gaussian filters: (1) filter centered at the flicker frequency *f* with full-width at half-maximum (FWHM) = 0.5 Hz; (2) filter centered at *f-1* Hz with FWHM = 2 Hz; (3) filter centered at *f+1* Hz with FWHM = 2 Hz. Data filtered at the flicker frequency are called “signal” (S), and data filtered at the neighboring frequencies are called “reference” (R). Second, temporally-filtered data from 500 to 2600 ms (relative to the mask onset) were used to compute channel covariance matrices (two R matrices and one S matrix). The first 500 ms contains visual evoked response to the onset of the flicker, which affects the quality of the spatial filter and thus were excluded from the analyses (Cohen and Gulbinaite, 2017). However, we did not exclude the data following imperative stimulus onset because congruent and incongruent trials do not differ in early sensory evoked potentials over occipital and parietal areas (Appelbaum et al., 2011). Third, generalized eigenvalue decomposition (Matlab function *eig*) on the covariance matrices R and S was used to construct spatial filters, where R is average of two reference (flicker-frequency neighboring) covariance matrices, and S is the covariance matrix from data that was temporally filtered at the stimulus frequency. The eigenvectors (column vectors with values representing electrode weights) were used to obtain RESS component time series (eigenvector multiplied by the original unfiltered single-trial time series). Although the first RESS component typically has highest SNR at the frequency of interest (i.e. EEG responses at the frequency that the spatial filter is designed to maximize), power spectra of RESS components were expressed in SNR units, and the RESS component with the highest SNR at the tagging frequency was automatically selected for subsequent analyses. Out of 162 RESS components (27 participants, 6 components per participant), the 1^st^ RESS component was selected for subsequent analyses in 156 cases, and the 2^nd^ RESS component in 6 cases. Thus, for each trial we analyzed two separate frequency-specific RESS component time series (one for target, and one for flankers). Topographical maps presented in Figure 2A were obtained from the filters by left-multiplying the eigenvector by the signal covariance matrix. Maps were normalized to allow averaging across participants.

**Figure 2.**
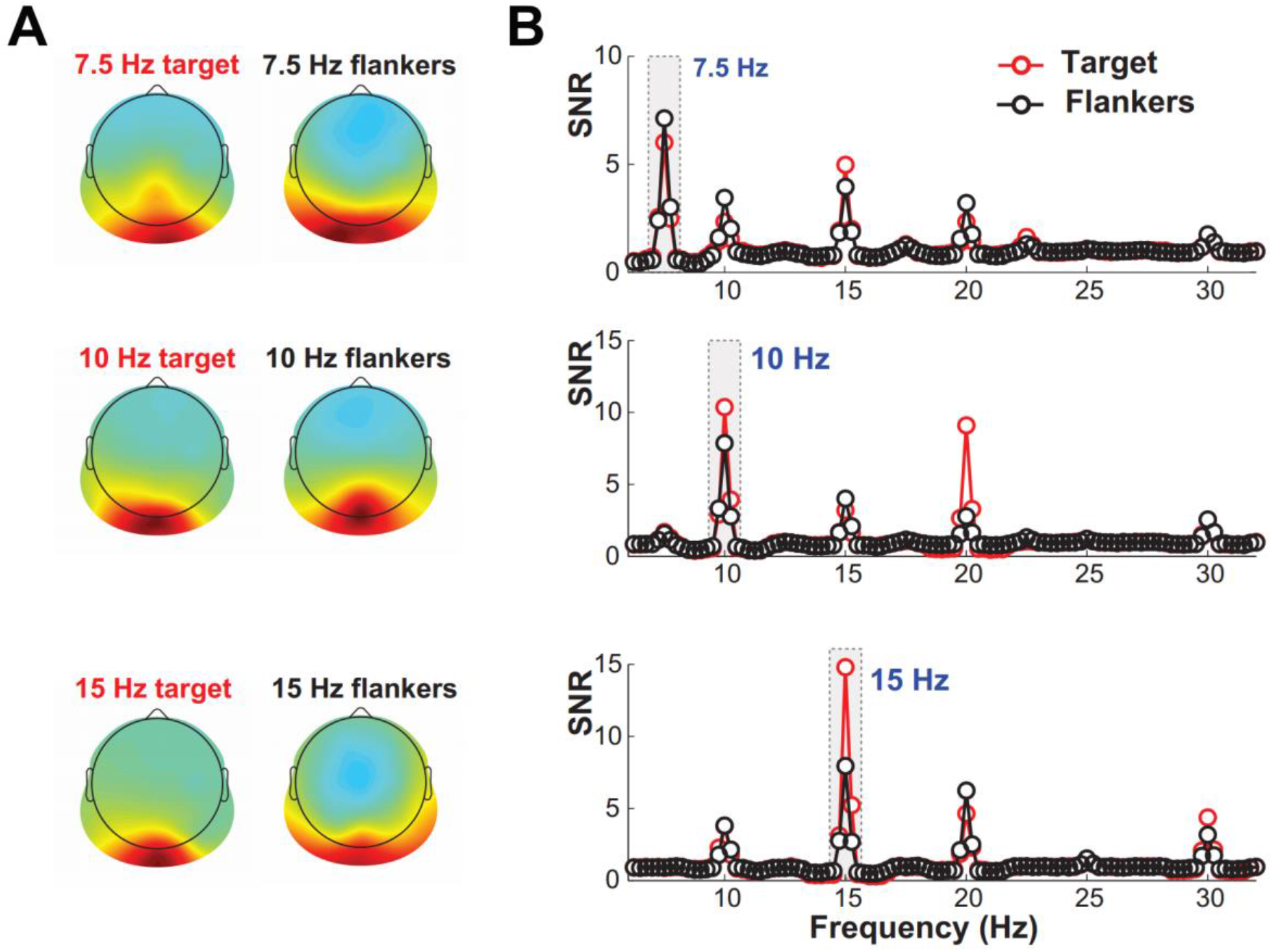
Topographical maps of frequency- and stimulus-specific SSVEP spatial filters and frequency spectra of RESS component time series. (A) RESS topographical maps for different tagging frequencies at target and flanker positions. Note that topographical distribution of SSVEPs elicited by the target is more centrally focused than that of the flankers. (B) Frequency spectra of RESS component time series, expressed in SNR (signal-to-noise ratio) units, highlights frequency-specificity of RESS SSVEPs. Note that 10 Hz target and 10 Hz flanker spectra are based on averaging 10T/xF and xT/10F conditions (where x is 7.5 or 15 Hz).

Although the electrode-level frequency domain representation of SSVEPs revealed peaks at the stimulus frequency (e.g. 10 Hz) and its harmonics (e.g. 20 Hz and 30 Hz), indicating well-documented nonlinearities in the response of the visual system (for a review, see Norcia et al., 2015), we focused our analyses on the fundamental frequencies (7.5. Hz, 10 Hz, and 15 Hz). As detailed in the Introduction section, we had specific a-priori hypotheses regarding alpha-band (10 Hz) vs. non-alpha (7.5. Hz and 15 Hz) flicker in terms of interactions with endogenous alpha oscillations implicated in the Eriksen flanker task (Fan et al., 2007; McDermott et al., 2017), and attentional processes in general (Jensen and Mazaheri, 2010; Klimesch, 2012). Although responses at higher harmonics and inter-modulatory frequencies reflect nonlinearities of the visual system, and have been used to study low-level visual processes (e.g. adaptation, symmetry, binocular rivalry; for a review see, Norcia et al., 2015), yet the physiological origin (retinal, subcortical, or cortical) has not been systematically investigated (Kim et al., 2011; Labecki et al., 2016).

### Theta-band (3-7 Hz) power

Frontal midline theta power, a well-established marker of response conflict task performance (Cavanagh and Frank, 2014; Cohen, 2014), was estimated using spatiotemporal filtering (procedure similar to that utilized for SSVEP source separation). This was done to optimize a theta component, which increases the accuracy of estimating the true neural theta activity (Cohen, 2017a). This is particularly important for the single-trial analyses. The data for the reference matrix was the broadband EEG data, and the data for the signal was the data temporally filtered around subject-specific condition-average theta-band peak frequency (FWHM = 3 Hz) which was defined in the following steps. First, we convolved Laplacian-transformed single-trial data from all electrodes with the complex Morlet wavelets, defined as: *e*^*i*2*πf_i_t*^*e-t*^2^/^(2σ^2^)^ (where *t* is time, *f_i_* is frequency which ranged from 2 to 30 Hz in 40 logarithmically spaced steps, and *σ* is the width of each frequency band defined as n/(2πfi), where *n* is a number of wavelet cycles that varied from 4 to 6 in logarithmically spaced steps). Second, we computed instantaneous power by taking the square of the complex convolution result, and normalizing power values by converting to decibel scale relative to the pre-stimulus time window (-500 – -200 ms, where 0 is the onset of the flickering mask). Third, given that theta-band (3-7 Hz) activity around response time (2300 – 2600 ms) was maximal at FCz and Cz electrodes (Fig. 3B left topoplot), the average TF map of these two electrodes was used to automatically find condition-average subject-specific theta peak frequency in post-stimulus window. The data temporally filtered around this peak frequency was used for constructing the spatiotemporal filter that maximized midfrontal theta signal.

**Figure 3.**
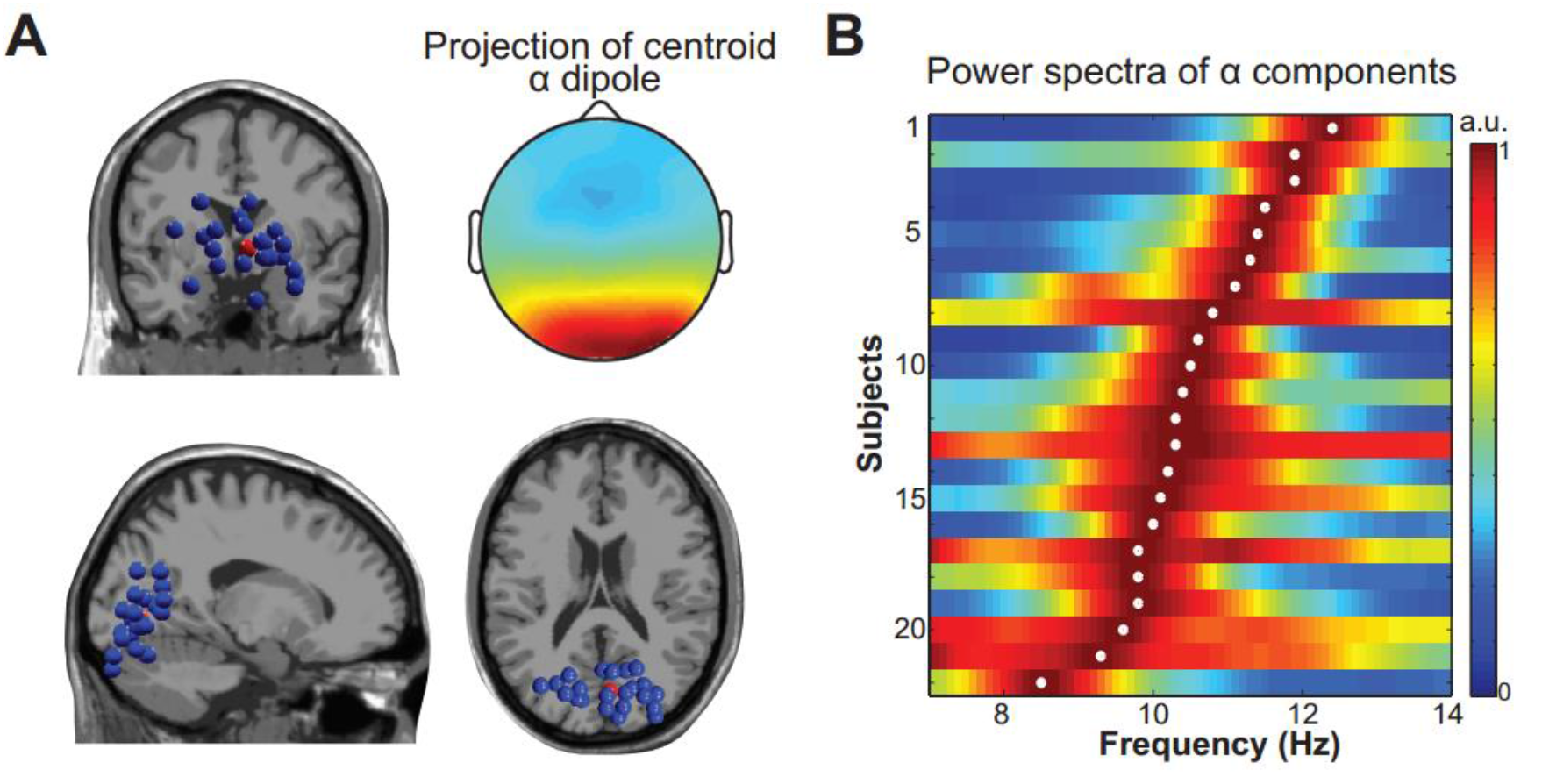
Occipital alpha sources. (A) Locations of equivalent dipoles for occipital alpha independent components (ICs). Each blue dot represents a participant. The red dot represents the centroid of the cluster based on all participant dipoles, the scalp projection of which is depicted in the top right. (B) Power spectra of alpha IC time series, normalized to the power of each participant’s alpha peak (for comparability across participants). Each row represents a participant, color corresponds to normalized spectral power, and the white dot denotes alpha peak frequency.

Data from 2000 to 2800 ms (relative to the mask onset) were used to compute two time-averaged covariance matrices (S and R) across all electrodes, which were used for generalized eigenvalue decomposition. The eigenvector with the largest eigenvalue was used as a spatial filter that, when multiplied by the raw EEG data (de Cheveigne and Arzounian, 2015), provided a component that maximizes theta-band activity. Forward model projections of the spatial filters were then visually inspected for characteristic midfrontal topography. For 26 out of 27 participants the 1^st^ eigenvector satisfied this criterion; and for 1 participant the 2^nd^ eigenvector was selected. Time-frequency representation of component time series was obtained using Morlet wavelets following the procedure identical to electrode-level data as described above.

### Occipital alpha-band source separation

In order to test our hypothesis that behavioral effects of 10 Hz flicker depend on the distance between 10 Hz and IAF, it was important to estimate occipital IAF as accurately as possible (Gulbinaite et al., 2017). For this, we determined occipital IAF using independent component analysis (ICA). We focused on occipital alpha, because for low-level visual stimuli (e.g. full-field flicker, pattern reversal of checkerboard and gratings) – similar to the stimuli used here – the effects of rhythmic stimulation are maximal in primary sensory cortices (Muller et al., 1997; Di Russo et al., 2007; Cottereau et al., 2011). Moreover, we determined IAF from the inter-trial interval, because IAF not only differs across brain regions, but also is state-dependent (task-related vs. resting-state IAF Haegens et al., 2014). Furthermore, there was no flicker during the inter-trial interval, so our estimation of occipital IAF was not biased by SSVEPs.

First, eye-movement artifact-free data were band-pass filtered at 5-15 Hz. Second, the ICA was performed on temporally-filtered data using only the pre-stimulus time window (-1000 to 0 ms, where 0 is the onset of the mask), obtaining 20 independent components (ICs). For each independent component, a single equivalent current dipole model was fitted using three-layer BEM template model based on the standard Montreal Neurological Institute's (MNI) brain template from the DIPFIT plug-in (DIPFIT toolbox; Oostenveld and Oostendorp, 2002). The occipital IC was selected based on proximity to occipital ROIs centered on Brodmann areas 17 and 18 (right-side MNI coordinates: -20 -70 50; left-side MNI coordinates: 20 -70 50; Haegens et al., 2014), with constraints that the selected equivalent dipole had less than 15% residual variance from the spherical forward-model scalp projection, and was located inside the model brain volume. The average residual variance of the dipole fit for the selected occipital ICs was 5.8% (SD = 4.37%). Finally, occipital IAF was estimated by taking the FFT of the IC time series in the -1000 – 0 ms window. The data was zero-padded to obtain 0.1 Hz frequency resolution. The absolute value of FFT coefficients was squared and averaged across trials. The individual alpha-peak frequency was determined as the peak in the range of 6-14 Hz. This frequency search window was selected based on reports that IAF ranges from 6-14 Hz (Bazanova and Vernon, 2014). For five participants, IAFs could not be determined due to small alpha peaks in the power spectrum that were indistinguishable from noise. Thus 22 participants were included in the single-trial analyses using linear mixed-effects models described further. All IAFs were above 8 Hz and below 13 Hz, thus there was no interaction with control flicker frequencies of 7.5 Hz and 15 Hz (Fig. 3B).

### Statistical analyses

#### Trial-average behavior

We tested the effect of flicker on average behavioral performance using repeated-measures ANOVAs. First, we kept the 4 tagging conditions separate, and entered mean RT and percentage error data in two separate repeated measures ANOVAs with factors trial congruency (congruent, incongruent) and tagging condition (7.5T/10F, 10T/7.5F, 10T/15F, and 15T/10F; the first number denotes target flicker frequency) as within-subject factors. Second, we evaluated a general effect of 10 Hz flicker in flanker vs. target positions by collapsing conditions, and submitting mean RTs and percentage error to another set of repeated-measures ANOVAs with factors trial congruency (congruent, incongruent) and tagging condition (10 Hz target, 10 Hz flankers).

#### Trial-average SSVEP analysis

SSVEP amplitude was calculated by performing single-trial FFTs of RESS component time series in a 500-2600 ms time window (relative to the mask onset). The first 500 ms were excluded to remove stimulus-evoked activity at the trial onset (Andersen et al., 2011). To obtain equally good resolution for all flicker frequencies (0.25 Hz), the exact time window for FFT was adjusted by zeropadding the data. Absolute value of FFT coefficients was averaged across trials and squared, and converted into SNR units to facilitate comparison across different flicker frequencies (Norcia et al., 2015). SNR was computed as the power at the flicker frequency of interest divided by the average power at the neighboring frequencies (+/-1 Hz, excluding +/-.5 Hz around the frequency of interest; e.g. 8.5-9.5 Hz and 10.5-11.5 Hz for 10 Hz flicker frequency).

To evaluate the amount of attentional modulation across different flicker frequencies, SNR values at the flicker frequency of interest were submitted to a two-way repeated measures ANOVA, with stimulus type (target, flankers), and frequency (7.5, 10 Hz paired with 7.5 Hz, 10 Hz paired with 15 Hz, and 15 Hz) as within-subject factors.

#### Trial-average theta-band power

Based on visual inspection of the subject-average theta component time-frequency plot, a window with the largest theta-band power (3-8 Hz, 2200-2800 ms; marked with a white rectangle in Fig. 5A bottom) was used to find participant- and condition-specific time-frequency peaks. An automatic peak finding procedure was adopted to capture individual differences in task-related brain activity. Average peak values (+-1 Hz, +-150 ms around the time-frequency peak) were entered in 2 (congruency) x 4 (tagging condition) repeated measures ANOVA.

#### Single-trial analyses

Single-trial analyses were performed using linear mixed-effects models (LMEs). LME models are extensions of standard regression models, and allow to simultaneously assess the influence of several predictors (i.e. fixed effects), while taking into account within-subject variability (i.e. random effects). The random effects are included in the model as so-called random intercepts and random slopes, which ensures that the observed effects are not driven by the data of one subject. Thus LMEs provide a more accurate and sensitive understanding of the patterns in the data.

Model fitting was implemented using the *lmer* package in R software (Bates et al., 2015). Singletrial RT was used as a dependent variable and was log-transformed to correct for the positive skew of RT distribution. In the reported models, fixed effects represent the general relationship between single-trial RTs and experimental factors, whereas random effects reflect subject-specific deviations from this general pattern. Specifically, random intercepts model the variability in RT of individual subject (some subjects may be fast, while others may be slow), whereas the random slopes model the variability in the influence of experimental factors on RTs (i.e. the effect of congruency on RTs may be stronger for some subjects than for the others).

The best-fitting model was selected using an iterative procedure. First, we fitted the base model which included only trial congruency as a fixed factor, and a random intercept term to account for subject-specific variability in the offset. Thereafter we gradually increased model complexity by adding additional fixed factors and their interactions, while ensuring the model’s goodness of fit by comparing Akaike information criterion (AIC), and selecting a more complex model if decrease in AIC was more than 2. The decrease in the Akaike Information Criterion (AIC) may be interpreted as evidence ratio. For example, a (more complex) model with a lower AIC is e^(AIC difference/2)^ times more likely to represent the data than the model with a higher AIC. We used an AIC threshold of 2, which means that a more complex model must be at least 2.7 times more likely to represent the data than the simpler model (Akaike, 1974). Thus, additional factors were included only if they explained a significant amount of variance (Baayen et al., 2008). We also validated the model by plotting residuals against the fitted values, and performed model criticism by removing 1.5% of the data (potential outliers). Plotting and estimation of p-values for each factor in the best-fitted model were performed using *sjPlot* package (Lüdecke, 2017).

The following factors were used as RT predictors: Trial congruency (2 levels), tagging condition (4 levels), single-trial theta power, single-trial SSVEP power at target and flanker tagging frequencies (expressed in SNR units using identical procedure to that of trial-average), distance between occipital IAF and 10 Hz (i.e. absolute difference). Single-trial theta power was defined from the time-frequency representation of the theta component (no baseline correction applied), by taking the average around theta power peak (+-1 Hz; +-150 ms) defined separately for each participant from condition-average theta component time-frequency plot (see Fig. 3A bottom). Single-trial SSVEP power at target and flanker tagging frequencies was defined by taking the FFT of respective RESS components in two time windows: Pre-stimulus (1400-2000 ms relative to the mask onset), and post-stimulus (2000-2600 ms). This choice was motivated by previous reports on alpha power modulations over brain regions representing ignored locations prior to and after stimulus onset (Handel et al., 2011; van Diepen et al., 2016). We reasoned that the effects of 10 Hz flicker might have a different effect on proactive vs. reactive attention control.

All numerical predictors (theta power, SSVEP amplitude for target and flanker stimuli) were normalized to a Gaussian distribution by ranking the data, scaling between -1 and 1, and taking the inverse hyperbolic tangent (also known as Fisher transform; Cohen, 2017b). Categorical factors were dummy-coded: “congruency” was coded as 0 or 1 (1 for congruent and 0 for incongruent trials), and “tagging condition” was coded 1 to 4 (10T/7.5F as 1, 10T/15F as 2, 7.5T/10F as 3, 15T/10F as 4).

## RESULTS

### Behavioral results: Trial-average analyses

Based on the findings that alpha power is increased over task irrelevant regions and has been interpreted to reflect inhibition of task-irrelevant information (Handel et al., 2011; Frey et al., 2015), and on a recent report on hemifield-specific entrainment of alpha-band oscillations using 10 Hz flicker (Spaak et al., 2014), we predicted 10 Hz flicker to have the following behavioral effects. We expected responses to be faster and more accurate when flankers flickered at 10 Hz, because inhibition of the distractors would facilitate processing of the target stimulus. On trials where the target stimulus flickered at 10 Hz, we expected responses to be slower and less accurate, because increase in alpha-band oscillatory power would be detrimental for processing in the task-relevant regions. A null result may indicate that (1) individual differences in IAF have to be taken into account or (2) alpha-band dynamics may reflect more global processing (e.g., hemifield-specific as found by Spaak et al., 2014) than spatially local processing as in our experiment (Fig. 1A).

A summary of the behavioral results is illustrated in Figure 4. Despite our novel experiment design with flickering stimuli and the circular arrangement of the flankers, we observed a typical behavioral pattern previously reported in the Eriksen flanker task (Wendt et al., 2007; Nigbur et al., 2012; Gulbinaite et al., 2014): Responses were faster (F(1,78) = 11.73, p = .002, η^2^ = .311) and more accurate (F(1,78) = 31.14, p < .001, η^2^ = .545) on congruent as compared to incongruent trials. This congruency effect was observed across all four tagging conditions, as indicated by the non-significant congruency by tagging condition interaction, both for RTs (F(3,78) = 0.01, p = .999, η^2^ = .001), and error rates (F(3,78) = 0.43, p = .734, η^2^ = .016). The main effect of tagging condition was not significant for error rates (F(3,78) = 0.71, p = .546, η^2^ = .027), but was significant for RTs (F(3,78) = 7.84, p < .001, η^2^ = .232). Follow-up Bonferroni-corrected comparisons revealed that participants on average responded the slowest when flankers flickered at 15 Hz (10T15F condition), and response speed in the latter condition significantly differed from 10T/7.5F (p = .036) and 15T/10F condition (p < .001), but not from 7.5T/10F condition (p = .095).

**Figure 4.**
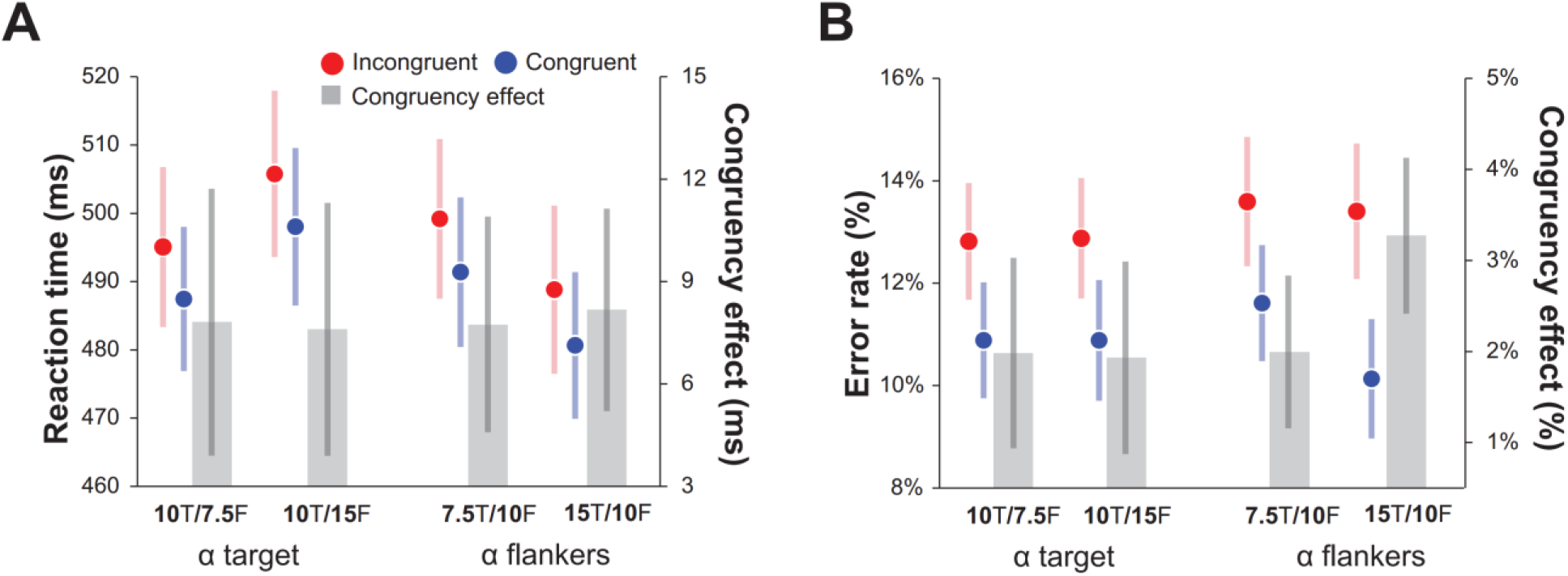
Behavioral performance. (A) Mean reaction time and (B) error rates on congruent vs. incongruent trials plotted as a function of condition. Gray barplots in each graph represent congruency effect (RT and error-rate difference between incongruent and congruent trials). Error bars denote standard error of the mean.

We next tested whether 10 Hz in the target vs. flanker positions differentially affected behavioral performance. As predicted based on the inhibitory role of alpha oscillations (Jensen and Mazaheri, 2010), participants responded significantly faster when flankers flickered at alpha (10 Hz) compared to non-alpha flanker flicker frequencies (F(1,26) = 8.27, p = .008, η^2^ = .241). However, follow-up analyses based on separate two-way ANOVAs revealed that this result was driven by significant differences in RTs between 15T/10F and 10T/15F conditions (F(1,26) = 40.40, p < .001, η^2^ = .608), whereas 7.5T/10F and 10T/7.5F conditions did not significantly differ (F(1,26) = 1.39, p = .249, η^2^ = .051).

In conclusion, average RT and accuracy analyses revealed that the combination of tagging frequencies rather than 10 Hz flicker at the target or flanker position had an effect on behavioral performance. Therefore, for the single-trial analyses we kept all four tagging conditions separate.

### Conflict-related theta-band power

Despite the presence of flicker, we observed a typical increase in theta-band power compared to the baseline period over the midfrontal electrodes (Fig. 5). In line with previous reports (Cohen and Cavanagh, 2011; Nigbur et al., 2012), the topographical distribution of stimulus-locked theta power peaked around FCz and Cz electrodes (Fig. 5B left). The forward model of the theta component obtained using source separation showed a spatial peak at FCz (Fig. 5B right) and an overall similar pattern of trial-averaged time-frequency dynamics (Fig. 5B). Theta power relative to the baseline period was increased more for incongruent than congruent trials (F(1,78) = 7.98, p = .009, η^2^ = .235), and this effect did not differ across tagging conditions (F(3,78) = 0.913, p = .439, η^2^ = .034). Theta power was most increased for 10T/15F, which was also the slowest RT condition. Post hoc Bonferroni-corrected comparisons revealed significant differences between 10T/7.5F and 10T/15F conditions (p = .004), as well as 10T/7.5F and 15T/10F conditions (p = .041).

**Figure 5.**
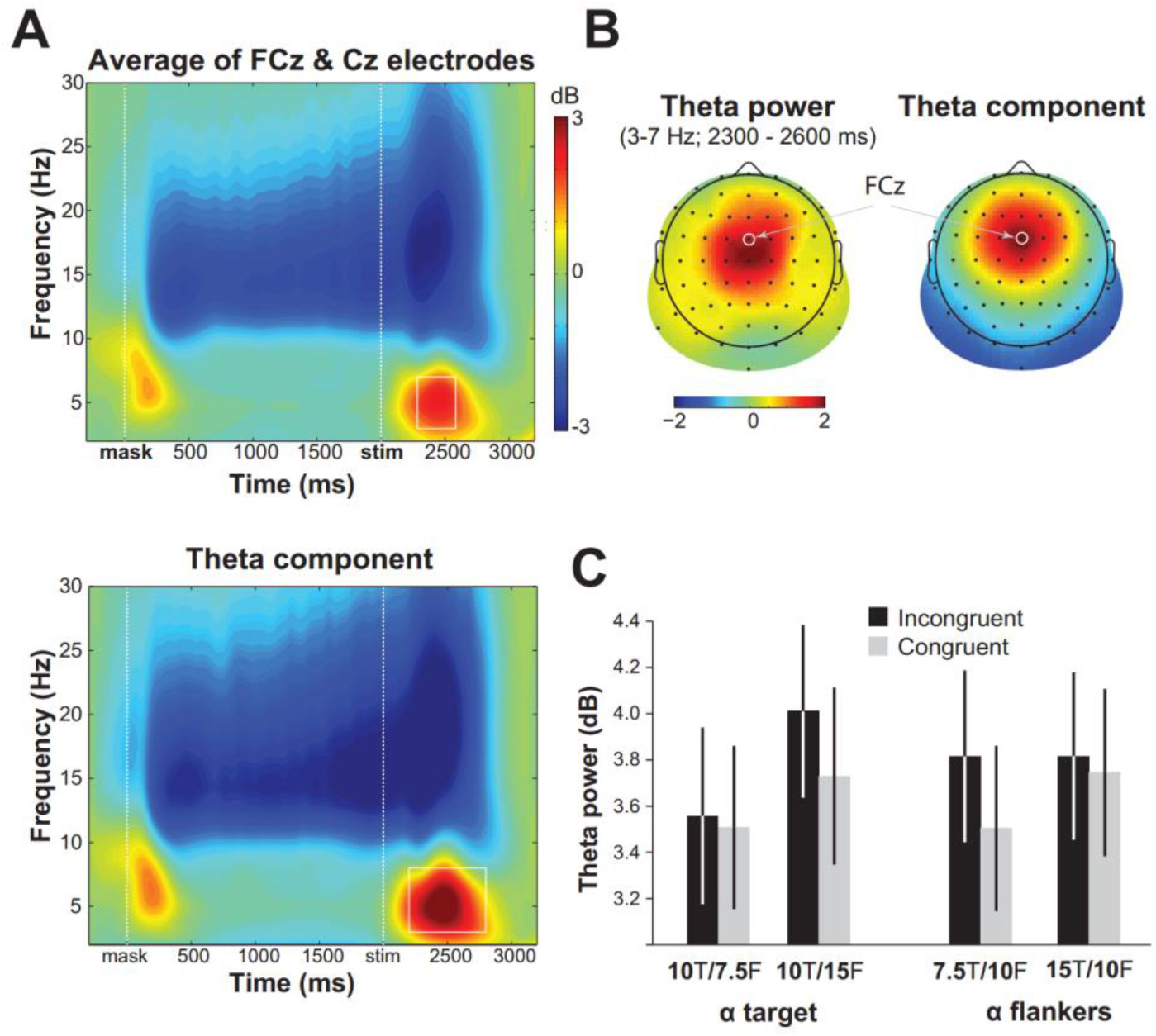
Conflict-related theta-band power. (A) Condition-average changes in power relative to the baseline period (-500 – -200 ms) at electrodes FCz and Cz (top panel), and theta component derived using source separation (bottom panel; see Methods section for details). (B) Theta-band (3–7 Hz) power distribution over the scalp in the time-frequency window indicated by the white square in panel A, and forward model of theta component. (C) Condition-specific changes in theta-band power (data taken from the theta component).

### 10 Hz flicker effects on SSVEP amplitude: Trial-average analysis

Although RESS spatial filters were designed to maximize brain responses to stimuli at specific spatial location (target vs. flankers) and specific tagging frequency (7.5 Hz, 10 Hz, or 15 Hz), frequency spectra of RESS component time series reveal responses at multiple harmonics of the stimulus frequency (2*f, 3f*; see Figure 2B). This result is consistent with previous reports showing partially spatially-overlapping cortical foci of the fundamental and higher harmonic responses (Pastor et al., 2007; Ales et al., 2012; Heinrichs-Graham and Wilson, 2012). Due to our a-priori theoretical motivation to focus on alpha-band flicker (see Introduction section), and conflict-related modulation of alpha-band power in the Eriksen flanker task (Fan et al., 2007; McDermott et al., 2017), we focused our analyses on the part of the SSVEP signal corresponding to the stimulus frequency (i.e. first harmonic, *f*) as opposed to higher harmonics (e.g. 2*f*, 3*f*). Here we report SSVEPs for different tagging frequencies (7.5 Hz, 10 Hz, and 15 Hz) at target and flanker positions during 500 to 2600 ms (relative to the mask onset) period. Note that 7.5 Hz target and 7.5 Hz flanker SSVEP results come from different blocks in the experiment.

Based on numerous studies using flicker for tagging dynamics of attention over space and time, we expected to observe higher SSVEP amplitude for target vs. flanker stimuli. Moreover, overall differences in SSVEP amplitude across tagging frequencies were predicted based on previously reported non-linearities in SSVEP response to wide-range flicker frequencies, with enhanced amplitudes, or resonance, at ~10 Hz (Regan, 1989; Herrmann, 2001) and 15 Hz (Pastor et al., 2003).

Consistent with the results of our previous study (Gulbinaite et al., 2014), SSVEP amplitude for attended stimulus (target) was on average higher than for ignored stimuli (flankers), as indicated by the main effect of stimulus type (F(1,26) = 6.20, p = .019, η^2^ = .193; Fig. 6). Significant stimulus type and flicker frequency interactions revealed that this attentional modulation was present not for all flicker frequencies (F(3,78) = 15.23, p < .001, η^2^ = .369). A pair-wise t-test revealed that attentional modulation was significant for 10 Hz tagging frequency when it was paired with 7.5 Hz (p = .024) and 15 Hz tagging frequency (p < .001), and no significant attentional modulation for 7.5 Hz tagging frequency and 10 Hz tagging frequency paired with 15 Hz. There was also a main effect of flicker frequency (F(3,78) = 14.71, p = .001, η^2^ = .361), such that SSVEP amplitude for 7.5 Hz flicker was significantly lower than for 10 and 15 Hz (p < .01).

**Figure 6.**
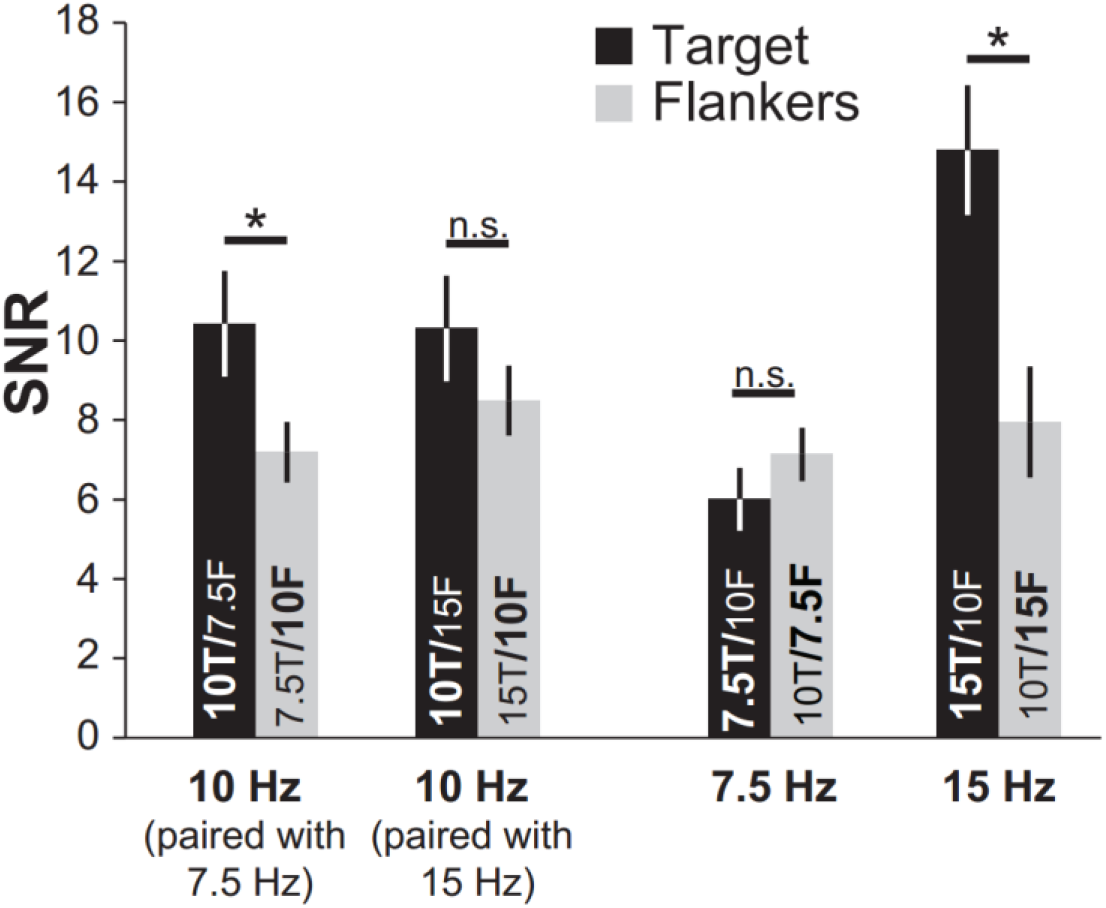
Attentional modulation of SSVEPs. Grand-average SSVEP amplitudes (determined from 500 to 2600 ms time window and expressed in SNR units) for each stimulus type (target and flankers) and each flicker frequency. The data plotted here is taken from the frequency spectra depicted in Figure 2, except that 10 Hz target flicker conditions are separated. Error bars denote standard error of the mean. * indicates p < .05, n.s. indicates p > .05.

### 10 Hz flicker effects: Single-trial analysis

To evaluate the effects of 10 Hz flicker on behavioral performance (reaction time) at the level of individual trials, we combined single-trial EEG measures (target and flanker SSVEPs, theta-band power) and experimental factors (IAF distance to 10 Hz, congruency, and condition) using linear mixed-effects models (LMEs). As noted in the Methods section, we used log-transformed single-trial RT as a dependent variable. Single-trial analyses were implemented into two steps. First, we derived a best-fitting mixed-effects regression model using an iterative model selection procedure. Second, we tested significance of fixed effects included in the best-fitting model.

We started with the baseline model, which included trial congruency as a fixed-effect factor, and a random intercept term to account for subject-specific variability in the RT offset. Our second model additionally included a fixed effect of condition (coded as “1” for 10T/7.5F, “2” for 10T/15F, “3” for 7.5T/10F, and “4” for 15T/10F), which significantly improved the model fit (decrease in AIC of 54.2; ⍰^2^(3) = 60.23, p < .001). We then tested whether the effect of condition and/or congruency varied across participants by including random factors of congruency and condition. Inclusion of condition as an additional random effect further improved the model fit (decrease in AIC of 20.5; ⍰^2^(9) = 38.43, p < .001). Further on, fixed and random effects of additional predictors were included only if they significantly improved model’s goodness of fit to the single-trial RT data. Table 1 summarizes fixed-effect factors that were gradually added to the model, and indicates each model’s goodness of fit in comparison to a simpler model (with respect to the fixed-effects structure). While results in Table 1 are based on a baseline random-effects structure (only a random intercept term), random slopes for each additional predictor were included whenever model comparison showed these to be necessary (full model selection procedure can be found in https://figshare.com/s/e6ad00be61a10bfe6a98).

**Table 1.** Illustration of stepwise best-fitting statistical model selection procedure. The first row represents the baseline model which includes the fixed effect of congruency and a random intercept per subject. In each subsequent row, additional fixed-effects factors and their interactions (denoted with colon sign) are gradually included. Increase in log-likelihood and decrease in AIC indicates an increase in goodness of fit obtained by adding an additional predictor in comparison to the simpler model in the row above. Note that all models presented in this table had only a random effect of intercept. alphaABS = distance between 10 Hz and individual alpha peak frequency (IAF); Target-SSVEP and Flankers-SSVEP = single-trial SSVEP amplitude in the post-stimulus time window (2000-2600 ms) for target and flanker stimuli respectively.

| Models | Log-lik.<br>increase | AIC<br>decrease |
| --- | --- | --- |
| 1. RT ~ congruency + (1\|Subj) | - | - |
| 2. ... + condition | 30.1 | 54.2 |
| 3. ... + alphaABS + congruency : alphaABS | 5.4 | 6.7 |
| 4. ... + Target-SSEP : condition + Flankers-SSEP : condition | 86.3 | 156.5 |
| 5. ... + theta power | 61.6 | 121.3 |

Specifications of the best-fitting model and statistics of the fixed-factors are reported in Figure 7A. The model explained 30.31% of variance in single-trial RTs ((calculated using piecewiseSEM package; Lefcheck, 2016) (as recommended by Nakagawa and Schielzeth, 2013)). Residuals of our best-fitting model followed a normal distribution. In the model criticism procedure, removing 1.5% of the data (potential outliers) did not compromise the model fit. Regression weights, their associated p-values, and confidence intervals are visually represented in Figure 7A. In the text below, we highlight the effects that are most relevant for understanding the role of alpha-band flicker on task performance, and its dependence on IAF.

**Figure 7.**
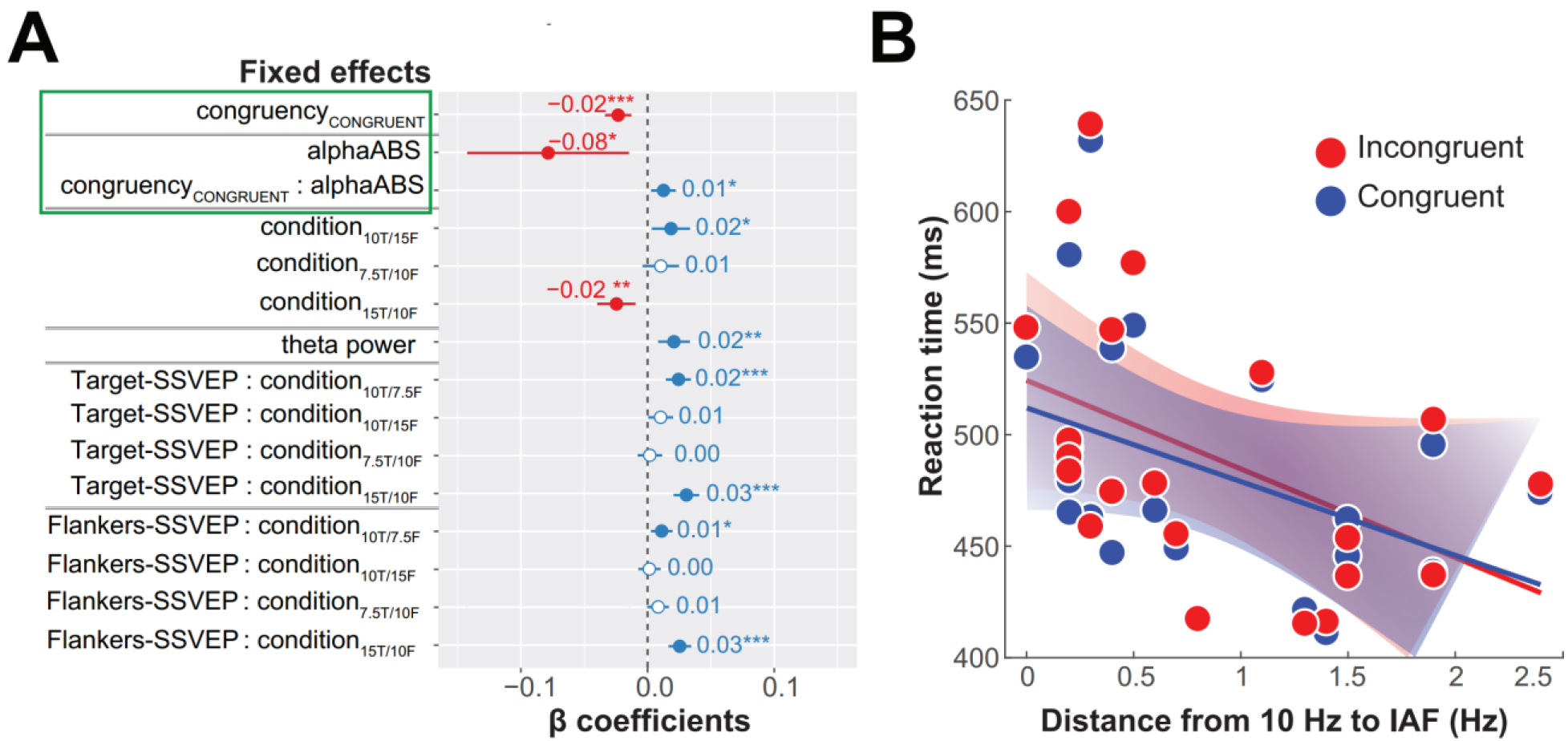
Single-trial analysis results. (A) Graphical representation of the best-fitting statistical model (fixed effects only), where: congruencyCONGRUENT indicates comparison between congruent and incongruent trials; congruencyCONGRUENT : alphaABS indicates differences in the effect of alphaABS on incongruent vs. congruent trials; condition10T/15F indicates comparison between conditions 10T/7.5F and 10T/15F; condition7.5T/10F indicates comparison between conditions 10T/7.5F and 7.5T/10F; condition15T/10F indicates comparison between conditions 10T/7.5F and 15T/10F. Error bars indicated 95% confidence intervals. Numbers denote fixed effects coefficients. Statistical significance of fixed effects coefficients is marked with asterisk symbols, where * is p < .05, ** is p < .01, *** is p < .001, and non-significant coefficients are marked with empty circles. (B) Graphical summary of fixed effects marked with a green rectangle in panel A, which illustrates that closer match between IAF and 10 Hz flicker resulted in overall slower RTs, and that this effect was stronger for incongruent trials (steeper regression line slope). Shaded areas represent 95% confidence intervals around the slope of regression line. Note that for LME modeling log-transformed RTs were used, however here mean RTs are left in original units for interpretation clarity.

The effect of condition corroborated the trial-average results. The regression weight for the 10T/15F condition was significantly positive, meaning that RTs were slower in 10T/15F relative to 10T/7.5F condition (⍰ = 0.02, (SE) = 0.008, t = 2.39, p = 0.026). The regression weight for 15T/10F condition was significantly negative, indicating faster RTs in 15T/10F vs. 10T/7.5F condition (⍰ = -0.025, (SE) = 0.008, t = -3.19, p = .004). Note that 10T/7.5F condition served as a reference condition.

The strongest predictor for log-RT was the absolute difference between 10 Hz flicker and individual alpha peak frequency: The closer a participant’s IAF to the stimulation frequency of 10 Hz, the slower the participant responded. This effect was stronger for incongruent than for congruent trials, as indicated by the interaction with congruency (regression weight for incongruent trials ⍰_alphaABS_ + ⍰_alphaABS:congruency_ = -0. 08 + 0*0.01, for congruent trials alphaABS ⍰_alphaABS_ + ⍰_alphaABS:congruency_ = -0. 08 + 1*0.01). Figure 7B illustrates the relationship between RT (expressed in milliseconds rather than log-transformed units for interpretation clarity) and IAF distance to 10 Hz, and illustrates that the congruency effect was larger for participants that had their IAF close to 10 Hz. To ensure that this effect was due to IAF distance to 10 Hz, rather than to IAF per se, we compared the best-fitting model to the model with IAF as a fixed factor. Goodness of fit significantly decreased (i.e. an increase in AIC of 8.2), indicating that behavioral effects of alpha flicker depended on the match between individual’s occipital alpha and flicker frequency.

We next inspected whether the response to flicker (i.e., SSVEP amplitudes) was associated with task performance. We found that larger single-trial SSVEP amplitudes were generally predictive of slower RTs. However, the effects of alpha flicker were not specific to 10 Hz flanker or 10 Hz target conditions. Flicker rate for target vs. flankers, on the other hand, showed a pattern: In conditions with faster target vs. flanker stimulus tagging frequency (i.e. 10T/7.5F Hz and 15T/10F Hz), reaction time was significantly slower when both target and flanker SSVEP amplitude was higher (significantly positive fixed effects coefficients for Target-SSVEP:cond_10T/7.5F_, Target-SSVEP:cond_15T/10F_, Flankers-SSVEP:cond_10T/7.5F_, Flankers-SSVEP:cond_15T/10F_). Trials with more frontal theta power were also associated with slower RTs (⍰ = 0.02, (SE) = 0.006, t = 3.25, p = .004), replicating previous results (e.g., Gulbinaite et al., 2014; Cohen & Cavanagh, 2011).

Modulations of alpha-band power have been suggested to support both proactive (anticipatory) and reactive (stimulus-related) attentional processes (Foxe and Snyder, 2011; van Diepen et al., 2016). Our experimental design allowed us to test the effect of alpha-band flicker on proactive and reactive attention by predicting RTs based on SSVEP amplitudes only during the mask period (anticipatory attention allocation to the central hash mark), or on SSVEP amplitudes only during the stimulus presentation. We therefore constructed a model that included SSVEP amplitude prior to the imperative stimulus (i.e. 1400-2000 ms window, where 2000 ms is the onset of the stimulus). The time window for pre-stimulus and post-stimulus SSVEP amplitude calculation was matched in length to make sure that any observed effects are not driven by SNR differences due to the FFT window size. Model comparison revealed that the model including post-stimulus (2000-2600 ms time window) SSVEP amplitudes was superior to the model including pre-stimulus (1400-2000) SSVEP amplitudes (decrease in AIC of 117.8). This result indicates that 10 Hz flicker interfered more with reactive than proactive selective attention.

Our finding of alpha-band-specific effects of flicker on behavior, which were maximal close to the natural frequency of endogenous alpha, is consistent with the idea that SSVEPs reflect entrainment of endogenous oscillations rather than a linear summation of transient event-related potentials (ERPs) generated to each stimulus flash (Capilla et al., 2011; Notbohm et al., 2016). Although the relationship between SSVEPs and ERPs is an active area of research (for a comprehensive discussion, see Norcia et al., 2015) beyond the scope of present paper, we note that our findings satisfy the main “Criteria for direct entrainment of brain oscillations through a periodic external drive” listed in the review paper by Thut et al. (2011).

## DISCUSSION

### Behavioral effects of rhythmic visual stimulation are frequency-specific

Although rhythmic visual stimulation (“flicker”) is primarily used to “tag” processing of low-level visual and high-level cognitive phenomena (for a review, see Norcia et al., 2015), there is some evidence that flicker may also entrain endogenous rhythms (Spaak et al., 2014), and thus can modulate cognitive processes supported by these rhythms (Williams, 2001; de Graaf et al., 2013). The goal of the present study was to provide empirical evidence for this idea by testing the effects of alpha-band flicker on processing of relevant and irrelevant information in a selective attention task. In contrast to the previous studies that assessed perceptual and cognitive effects of flicker following alpha flicker offset (“offline” effects), we tested alpha flicker effects during stimulus processing (“online” effects).

Based on in vivo and in vitro studies, the prerequisite for influencing the brain’s rhythmic activity through rhythmic stimulation is a frequency match between the two (Reato et al., 2013). Frequencies that do not match the endogenous rhythm can still entrain endogenous oscillations provided the intensity of stimulation is sufficiently high (Reato et al., 2013; Notbohm et al., 2016). So far, studies attempting to modulate alpha oscillations via rhythmic visual stimulation did not incorporate IAF, nor did they consider that the task-related IAF may differ from the resting-state IAF (Haegens et al., 2014). We hypothesized that 10 Hz flicker effects on performance in the selective attention task will be maximal when the frequency of the applied rhythm matches that of the task-related alpha-band oscillations.

We used the Eriksen flanker task (Gulbinaite et al., 2014), with target and flankers flickering within (10 Hz) or outside the alpha band (7.5 Hz and 15 Hz). The flicker frequencies outside the classical alpha-band range (8-12 Hz) are necessary to demonstrate that flicker effects are specific to the alpha band, which was rarely done previously (but see de Graaf et al., 2013). We found that the closer the match between occipital IAF (estimated from the inter-trial interval data) and 10 Hz flicker, the slower a given participant reacted and the larger was the flanker interference effect (RT difference between incongruent and congruent trials). In the context of previous reports on stronger decrease of alpha-band power on incongruent relative to congruent trials (Fan et al., 2007; McDermott et al., 2017), the present finding of differential effects of 10 Hz flicker on incongruent vs. congruent trials points to an interesting possibility: 10 Hz flicker effects on endogenous alpha-band networks supporting selective attention are stronger when the network is itself more active. However, further research, elucidating on the precise mechanism of this interaction, is needed.

Behavioral effects of rhythmic stimulation on perception and attention that are specific to the alpha band have rarely been investigated (de Graaf et al., 2013; Shapiro et al., 2017). However, close inspection of several studies that employed a range of flicker frequencies suggests that alpha flicker is a special case. In a lateralized spatial attention task using tagging frequencies that ranged from 8 to 23 Hz, Toffanin and colleagues found that performance accuracy was significantly decreased when the stimulus background flickered at 9.5 Hz as compared to the other flicker frequencies (Toffanin et al., 2009). In another selective attention task, target detection accuracy was numerically smaller for flicker frequencies ~10 Hz (Ding et al., 2006). The attentional blink phenomenon is most pronounced when the stimulus stream is in the alpha and low beta frequency range (Shapiro et al., 2017). Together, these findings suggest that even on a group-level, alpha flicker effects on performance can be observed. However, this effect, as we have shown here, may be difficult to uncover if the relationship between exogenous rhythm (flicker) and endogenous rhythms (neural oscillations) is not taken into consideration.

### Global vs. local effects of 10-Hz flicker

Based on the well-established inhibitory role of alpha oscillations (Rihs et al., 2007; Jensen and Mazaheri, 2010; Klimesch, 2012; Samaha et al., 2016), and evidence for hemifield-specific entrainment of alpha-band oscillations using 10 Hz flicker (Spaak et al., 2014), we hypothesized that stimulus processing in the Eriksen flanker task will be impaired in parts of the visual field flickering in alpha (i.e. 10 Hz). However, the single-trial analyses combining behavioral and EEG data revealed that flicker entrainment effects were not specific to target or flanker positions. Although participants responded significantly faster when flankers flickered at alpha vs. non-alpha frequencies, this RT advantage was present only for 15T/10F as compared to 10T/15F condition (replicating the results of our previous study where we used 10 Hz and 12.5 Hz tagging frequencies; Gulbinaite et al., 2014), but not for 10T/7.5F Hz vs. 7.5T/10F.

There are two non-mutually exclusive potential explanations for the finding of global rather than local alpha-flicker effects. First, a neural explanation that takes into account the effects of flicker at the network level (Ding et al., 2006; Srinivasan et al., 2006; Srinivasan et al., 2007; Lithari et al., 2016); second, a cognitive explanation that takes into account the structure of the task.

Although low-level flickering visual stimuli (e.g. checkerboards and gratings) primarily entrain activity in visual cortex (Muller et al., 1997; Di Russo et al., 2007; Cottereau et al., 2011), it has been shown that flicker can also modulate activity in larger networks that extend beyond early visual cortex (Ding et al., 2006; Srinivasan et al., 2006; Srinivasan et al., 2007). For example, SSVEP amplitude in frontal areas is increased for theta-band (4-8 Hz; Mentis et al., 1997; Srinivasan et al., 2007) and beta-band flicker (~25 Hz; Pastor et al., 2003; Pastor et al., 2007). Source-estimation studies of SSVEPs also revealed distributed sources over occipital, parietal, and frontal areas associated with different flicker frequencies (Muller et al., 1997; Pastor et al., 2003; Srinivasan et al., 2006; Di Russo et al., 2007; Pastor et al., 2007; Kim et al., 2011; Heinrichs-Graham and Wilson, 2012). Thus, one explanation for the global alpha-flicker effects observed in our task is that alpha-band flicker may have interfered not only with selective attention, but with the functioning of multiple networks operating in alpha band (Sadaghiani and Kleinschmidt, 2016). Indeed, it is possible that the reason why the 10 Hz flicker effects were not spatially-specific is that the flicker, though retinotopically restricted, entrained large-scale alpha brain networks that spread to other retinotopic positions. Having the alpha flicker only in one hemifield may minimize this large-scale entrainment, which could explain why Spaak et al. (2014) observed flicker effects that were spatially-specific (at the level of visual hemifields).

In the Eriksen flanker task, spatial attention is proactively directed to the centrally presented target stimulus, whereas flankers draw attention due to target-flanker feature similarity. This instantiates reactive attentional control – engagement of both spatial and feature-based attention to minimize the influence of incongruent flankers. Importantly, although both spatial and feature-based attention are supported by alpha oscillations (van Diepen et al., 2016; Vissers et al., 2016), feature-based attention operates in a spatially non-specific manner (Serences and Boynton, 2007; Andersen et al., 2008). Thus, it is conceivable that the global effects of alpha flicker we observed here reflect interference with feature-based attention. This line of reasoning is further supported by our finding that trial-by-trial RTs were more strongly predicted by EEG flicker responses during stimulus processing (feature-based and spatial attention) than during stimulus anticipation. Hash marks served as placeholders which allow to filter out flanker locations and to prepare for the upcoming stimulus using spatial attention, but feature-based attention could not be efficiently engaged before the onset of the target and flanker letters.

### Multilevel analyses for uncovering brain-behavior relationship within and across individuals

Here we used a combination of two analysis approaches that allowed us to uncover theoretically relevant patterns of results that might otherwise be inaccessible when using only subject- and trial-average approaches. The first was multivariate source separation, which we used to define an optimal spatial filter to maximize the EEG response to flicker (Cohen and Gulbinaite, 2017), and to define a spatial filter that maximized theta-band activity (Cohen, 2017a). Such spatial filters provide components that, relative to selecting a single electrode, increase single-trial signal-to-noise ratio and more accurately reconstruct source time courses.

The second analysis approach was linear mixed effect models (Baayen et al., 2008), which allowed us to apply a formal model comparison approach to a dataset with multiple levels of variance, including cross-subject factors such as IAF, and within-subject factors such as single-trial SSVEP amplitude and theta power. We believe that the combination of EEG source separation and mixed-effects modeling provides a promising approach to uncover nuanced relationships between brain activity and behavior.

## Conclusions

Despite the long history of rhythmic visual stimulation in human electrophysiology research (Berger, 1929; Adrian and Matthews, 1934), the question of whether flicker can directly modulate perceptual and cognitive processes via entrainment of endogenous brain rhythms remains unresolved. By using a selective attention task, we show that whether and how external rhythmic stimulation affects brain function depends on the interaction between endogenous brain rhythms and externally-driven oscillations. The results of this study also demonstrate the importance of choosing the appropriate tagging frequency based on the inherent speed of the cognitive process of interest.

## Acknowledgements

This research was funded by the ERC-CoG P-CYCLES (N°614244) awarded to Rufin VanRullen. Michael X Cohen is funded by the ERC-StG THETA 2.0 (N°638589).

